# Aβ initiates brain hypometabolism and network dysfunction via NOX2 activation: a potential onset mechanism of Alzheimer’s disease

**DOI:** 10.1101/2020.08.12.248492

**Authors:** Anton Malkov, Irina Popova, Anton Ivanov, Sung-Soo Jang, Seo Yeon Yoon, Alexander Osypov, Yadong Huang, Yuri Zilberter, Misha Zilberter

## Abstract

A predominant trigger and driver of sporadic Alzheimer’s disease (AD) is the synergy of brain oxidative stress and glucose hypometabolism starting at early preclinical stages. Oxidative stress damages macromolecules, while glucose hypometabolism impairs cellular energy supply and antioxidant defense. However, the exact cause of AD-associated glucose hypometabolism and its network consequences has remained unknown. Here we report NADPH oxidase 2 (NOX2) activation as the main initiating mechanism behind Aβ_1-42_-related glucose hypometabolism and network dysfunction. We utilize a combination of electrophysiology with real-time recordings of metabolic transients both *ex*- and *in-vivo* to show that Aβ_1-42_ induces oxidative stress and acutely reduces cellular glucose consumption followed by long-lasting network hyperactivity and abnormalities in the animal behavioral profile. Critically, all of these pathological changes were prevented by the novel bioavailable NOX2 antagonist GSK2795039. Our data provide the first direct experimental evidence for causes and consequences of AD-related brain glucose hypometabolism, and suggest that targeting NOX2-mediated oxidative stress is a promising approach to both the prevention and treatment of AD.

**Single sentence summary:** Beta-amyloid induces brain hypometabolism, network hyperactivity, and behavioral changes via NADPH oxidase-mediated oxidative stress, suggesting a novel therapeutic target for Alzheimer’s disease treatment.

## Introduction

The pathological processes that drive AD may begin several decades before the first clinical symptoms manifest^1–3^. Meanwhile, numerous human studies suggest a causal upstream role for Aβ in AD initiation^3–5^, and recent progress in AD biomarkers has revealed that β-amyloidosis is one of the earliest signs of AD pathogenesis, detectable at preclinical stages of the disease^1–3^. Interestingly, at this stage patients are still cognitively unimpaired but may demonstrate variable neuropsychiatric symptoms (e.g., apathy, depression, agitation/aggression, anxiety, and irritability)^6,7^, the emergence of which may correlate with amyloidosis^8^. Although Aβ toxicity alone may not be sufficient to cause cognitive deterioration, it likely triggers downstream pathological changes (i.e., tauopathy and neurodegeneration) that ultimately lead to cognitive decline at later stages^3,9^.

Early Aβ pathology is often detected concurrently with glucose hypometabolism^10–12,13^, another pre-symptomatic marker of AD that is also an accurate predictor of progression to AD in mild cognitive impairment (MCI) patients^10,14,15^. Since glucose utilization underlies vital brain functions such as energy supply and antioxidant defense^16^, it is not surprising that disturbances in glucose metabolism can lead to a chain of harmful consequences, and thus likely represent a major underlying cause of disease initiation and progression^12^. However, until now the exact causes and consequences of AD-associated glucose hypometabolism have been unknown, hampering the search for effective treatment.

Multiple studies have shown that oligomeric Aβ induces brain oxidative stress^12,17^, *and* in different cell types Aβ_1-42_ induces oxidative stress^18^, largely via activation of NADPH oxidase (NOX)^19–21^. NOX is the only known enzyme with the primary function of generating ROS^22^. NOX family of enzymes are transmembrane carriers that transport an electron from cytosolic NADPH to reduce oxygen to superoxide anion. There are seven isoforms of NOX with NOX1, NOX2 and NOX4 expressed in multiple brain regions including the cerebral cortex, hippocampus, cerebellum, hypothalamus, midbrain and/or striatum^23^, with NOX2 the dominant form expressed by microglia, neurons, and astrocytes^23–25^. Several lines of evidence including postmortem analyses of AD patients’ cerebral cortices indicate that oxidative stress -- particularly resulting from NOX2 activation -- plays a significant role in the development of AD^20,23,25–27^. Close relationship between the levels of Aβ and NOX2 activity has also been well documented in multiple studies^19,28–33^.

We show that NOX activation by oligomeric Aβ_1-42_ results in pathological changes in brain glucose consumption, hippocampal network hyperactivity, and neuropsychiatric-like disturbances in mouse behavior. All observed abnormalities were prevented by blocking NOX2, suggesting NOX may be a major molecular mechanism behind AD initiation and progression. Our results point to early intervention in NOX-induced oxidative stress as a potential effective approach to AD prevention and treatment.

## Results

### Aβ_1-42_ disrupts network glucose utilization both in brain slices and in vivo

Fibrillar Aβ_1-42_ (400nM) applied to hippocampal slices reduced network activity-driven glucose uptake nearly by half (Figure 1A, Table 1.1), an effect paralleled by and correlated with decreased release of lactate (Table 1.2, Figure S1A-C). To confirm our results *in vivo*, we injected Aβ_1-42_ i.c.v. in anesthetized mice; following Aβ_1-42_ injection, extracellular glucose transients in response to synaptic stimulation changed more than two-fold (Figure 1B, Table 1.3), indicating a reduction in glucose uptake of similar magnitude to what we observed in brain slices. In the living brain, extracellular glucose is the product of interplay between glucose supply from blood and cellular glucose uptake. Network activation leads to an increase in glucose uptake but the blood vessels rapidly dilate and the supply of glucose is upregulated to compensate, leading to an actual net increase of extracellular glucose as has been demonstrated previously^34,35^. Since Aβ did not affect glucose delivery -- network local field potential (LFP) response to stimulation did not change significantly (Figure 1B, bottom right; Table 1.4), nor did the baseline glucose concentration (not shown) -- a doubling in extracellular glucose transient amplitude represents reduced glucose uptake. Aβ_1-42_-induced glucose hypometabolism was underlain by decreased glycolysis, seen as reduced amplitudes of NAD(P)H fluorescence “overshoot” (Figure 1C, Table 1.5) together with an increase of oxidation (“dip”) amplitude^36^ (Figure 1C, Table 1.6). This was paralleled by a decrease in FAD fluorescence “undershoot” phase (Table 1.7, Figure S1F), further confirming disrupted glycolysis^36^. Aβ_1-42_ also induced a significant increase in activity-driven oxygen consumption (Figure 1D, Table 1.8) that positively correlated with changes in NAD(P)H oxidation (Figure S1D), an association that likely indicates upregulated mitochondrial respiration to compensate for reduced glycolysis^37,38^.

**Figure 1.**
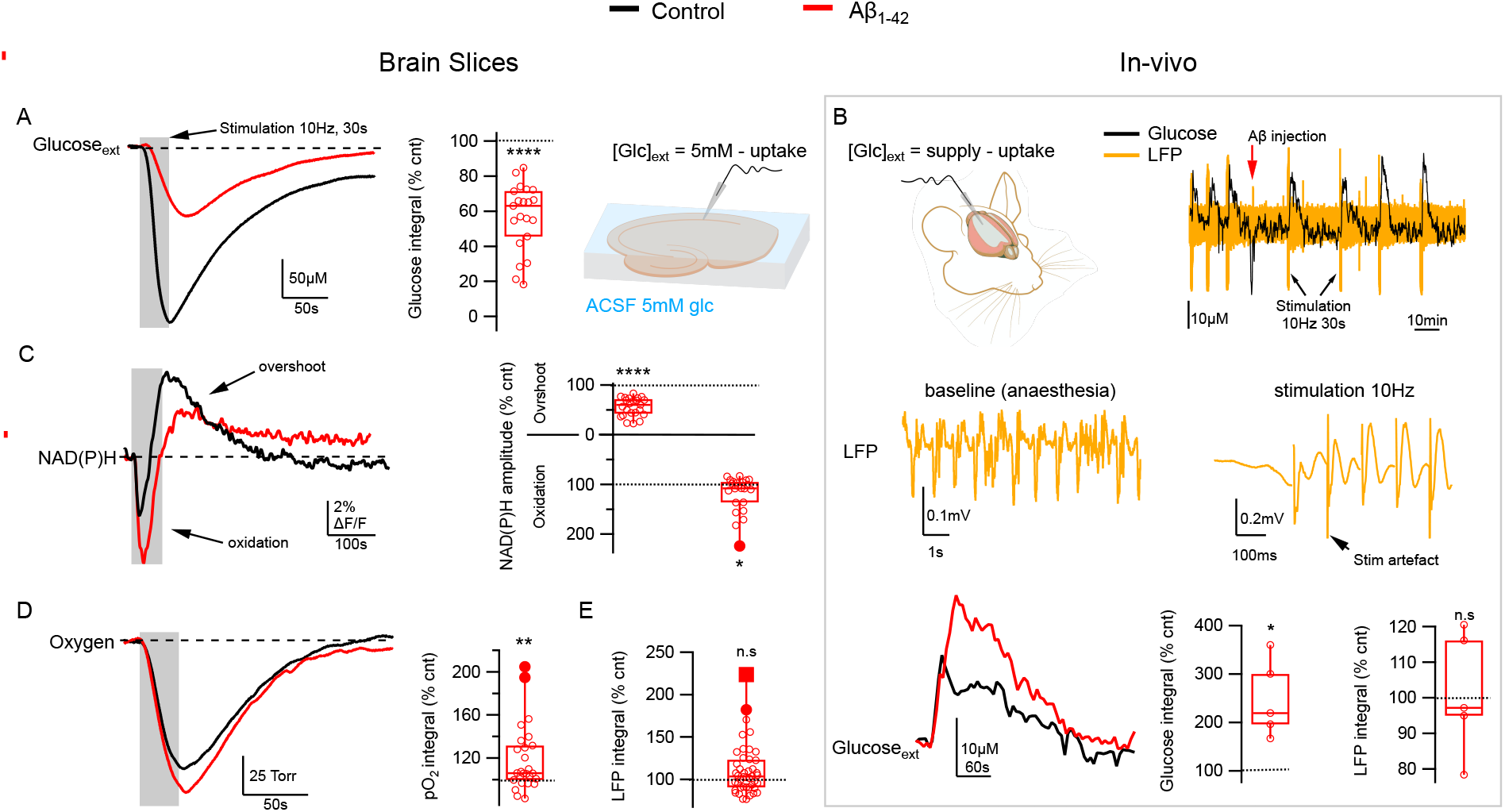
Aβ_1-42_ inhibits network glucose utilization. **A.** In brain slices, 40 minutes of fibrillar Aβ_1-42_ application reduces network activity-driven glucose uptake. *Left*, example traces from a single experiment showing the extracellular glucose transients in CA1 pyramidal cell layer in response to a 10Hz, 30s stimulation of Schaffer collaterals (grey) in control (black) and following 40min of Aβ_1-42_ application (red). Considering the constant 5mM glucose supply from the perfusate, the drop in the transient amplitude indicates reduced uptake. *Middle*, summary graph showing glucose transient integral values normalized to controls. **B.** In anesthetized mice, i.c.v. injection of fibrillar Aβ_1-42_ results in a rapid change in glucose uptake. *Top left*, a schematic depicting the relationship between the dynamic extracellular glucose supply from the blood (which increases following synaptic activation) and network glucose uptake. *Top right*: example traces from a single experiment showing both local field potential (LFP, orange) and extracellular glucose (black) recordings from the CA1 region before and after Aβ_1-42_ injection. *Middle row*: in detail of LFP recording showing characteristic anesthesia-induced oscillations (left) at baseline and the response to synaptic stimulation (right). *Bottom row*, average stimulation-induced glucose transients from a single experiment in control (black) and following i.c.v. Aβ_1-42_ injection (red). *Middle*, summary graph showing glucose transient integral values normalized to controls; *right*, a summary of LFP integrals showing that stimulation response did not change significantly following Aβ_1-42_ injection; This suggests that the activity-induced increase in glucose supply from the blood did not change, and therefore the apparent increase in glucose transients following Aβ_1-42_ injection indicates reduced uptake. **C.** Aβ_1-42_ inhibits activity-driven glycolysis. *Left*, NAD(P)H autofluorescence traces from a single experiment in control (black) and following 40min of Aβ_1-42_ application (red). *Right*, summary values of transient amplitudes both for the “overshoot” (glycolysis-related) and “oxidation” phases of the signal. **D.** Aβ_1-42_ increases oxygen consumption: sample pO_2_ traces from a single experiment showing a transient decrease of tissue oxygen levels in control (black) and following 40min of Aβ_1-42_ application (red) and a summary plot of normalized pO_2_ integral values. **E.** Aβ_1-42_ does not significantly change the strength of network response to synaptic stimulation: a summary plot of normalized LFP train integral values. *p <0.05, **p<0.01, ***p<0.001, ****p<0.0001

Meanwhile, none of the above disruptions could be attributed to changes in network activity, as the stimulus train LFP integral did not change (Figure 1E, Table 1.9), nor was there any significant correlation between observed metabolic changes and changes in network response (Figure S1E).

To test whether observed glucose hypometabolism could instead be due to reduced glucose transport as suggested previously^39–41^, we repeated NAD(P)H and FAD fluorescence recordings after increasing ACSF glucose concentration from 5mM to 10mM; if Aβ exerted its effect by inhibiting the passive glucose flux, then increasing extracellular glucose would serve to at least partially normalize that effect. However, doubling the glucose concentration in ACSF had no restorative effect on neither the disrupted NADH(P)H nor FAD transients (Figure S1F), suggesting Aβ - induced glucose uptake reduction is indeed underlain by reduced glycolysis and not by impaired glucose transport.

### Aβ_1-42_ toxicity is mediated by NOX2

Aβ_1-42_-induced changes in glucose utilization were strikingly similar to those we previously observed following seizures that were preceded by a spike in extracellular H_2_O_2_^42^. In that context, the likely source of the H_2_O_2_ and actual trigger of seizures is the reactive oxygen species (ROS)-generating NOX, specifically NOX2^43^. Aβ_1-42_ has been reported to induce oxidative stress in multiple cell types by activating NOX^18–21^. In our hands, when NOX2 was inhibited by the selective antagonist GSK2795039, Aβ_1-42_ failed to disrupt glucose utilization in slices (Figure 2A, Table 1.10) or *in vivo* (Figure 2B, Table 1.11). NOX2 inhibition also prevented Aβ_1-42_-induced modification of glycolysis (Figure 2C, Table 1.13-14) and oxygen consumption (Figure 2D, Table 1.15). Meanwhile, network activity response did not increase significantly following GSK2795039 or subsequent Aβ_1-42_ application (Figure 2E, Table 1.16), suggesting that positive changes in network response could not be responsible for the normalizing GSK2795039 effect. Finally, we repeated our experiments in slices from NOX2-deficient (Cybb^tm1din^/J) mice. Aβ_1-42_ failed to exert any detrimental effect in any of the observed parameters (Figure 2F, Table 1.17-20), confirming that Aβ_1-42_ inhibits glucose metabolism specifically via NOX2 activation-induced oxidative stress.

**Figure 2.**
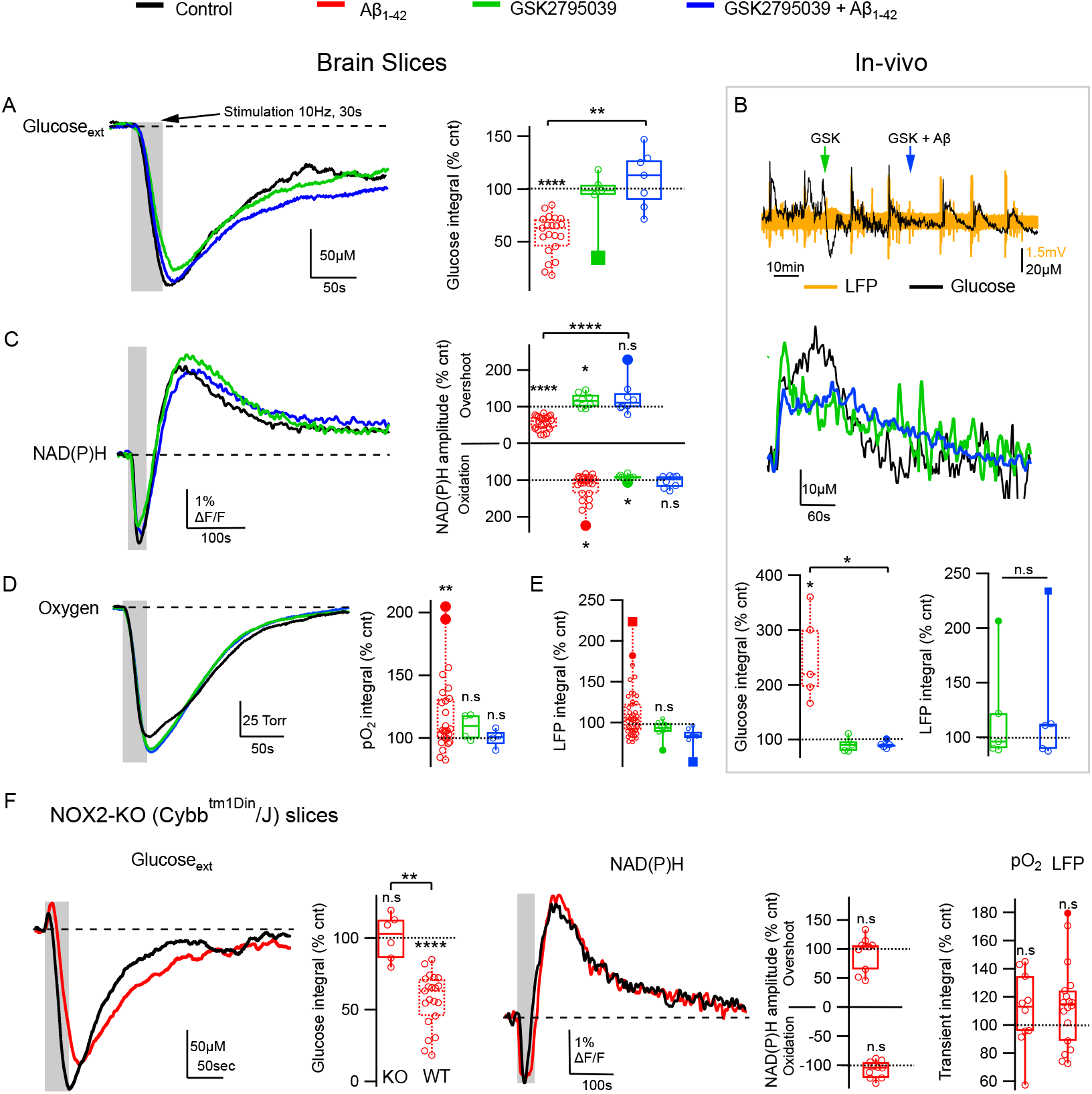
NOX2 is critical for Aβ_1-42_-induced disruption of network glucose utilization. In brain slices, application of NOX2 antagonist GSK2795039 prevents the reduction in network activity-driven glucose uptake caused by Aβ_1-42_: example traces from a single experiment showing the extracellular glucose transients in CA1 pyramidal cell layer in response to a 10Hz, 30s stimulation of Schaffer collaterals (grey) in control (black), after wash-in of GSK2795039 (green), and following 40min of GSK2795039 + Aβ_1-42_ application (blue); summary graph showing corresponding glucose transient integral values normalized to controls with those of Aβ_1-42_ alone for comparison (red). **B.** In anesthetized mice, intraventricular injection of GSK2795039 + Aβ_1-42_ (blue) preceded by GSK2795039 (green) fails to elicit any changes in glucose uptake. *Top*, example traces from a single experiment showing both local field potential (LFP, orange) and extracellular glucose (black) recordings from the CA1 region. *Middle*, average stimulation-induced glucose transients from a single experiment in control (black), following i.c.v. GSK2795039 injection (green) and subsequent injection of GSK2795039 + Aβ_1-42_ (blue). *Bottom left*, the summary graph showing corresponding glucose transient integral values together with those of Aβ_1-42_ alone for comparison (red). Bottom right, summary of LFP integral values showing no significant change following either injection. **C.** GSK2795039 prevents Aβ_1-42_ disruption of activity-driven glycolysis. *Left*, NAD(P)H autofluorescence traces from a single experiment in control (black) after washin of GSK2795039 (green), and following 40min of GSK2795039 + Aβ_1-42_ application (blue); *Right*, summary values of transient amplitudes both for the “overshoot” (glycolysis-related) and “oxidation” phases of the signal. **D.** GSK2795039 blocks Aβ_1-42_ induced increase in oxygen consumption: sample pO_2_ traces from a single experiment showing a transient decrease of tissue oxygen levels in control (black) after wash-in of GSK2795039 (green), and following 40min of GSK2795039 + Aβ_1-42_ application (blue); the summary plot of normalized pO_2_ integral values. **E.** GSK2795039 does not significantly modify network response to stimulation (green), while under Aβ+GSK2795039, the response decreases (blue); summary plot of stimulation train LFP integral values with Aβ-only values for comparison (dotted red). **F.** Aβ_1-42_ application has no effect in NOX2-deficient mouse slices: sample traces and summary of glucose (left) and NAD(P)H fluorescence transients (center), as well as summary of changes in oxygen consumption and LFP integrals (left). *p <0.05, **p<0.01, ***p<0.001, ****p<0.0001.

We further examined the role of NOX2 in oxidative stress induced by Aβ_1-42_ through additional biochemical analysis. Brain slices exposed to Aβ_1-42_ exhibited significantly increased oxidative stress seen as higher levels of malondialdehyde (MDA), a by-product of lipid peroxidation and a standard oxidative stress marker (Table 1.21, Figure S2). The Aβ_1-42_-induced increase in MDA levels was prevented by NOX2 inhibition by GSK2795039 (Table 1.22, Figure S2) and was also absent in Cybb^tm1dιn^/J mouse slices (Table 1.23, Figure S2), confirming that activated NOX2 is the primary source of overall acute oxidative stress induced by Aβ_1-42_.

We also tested whether alternative pharmacological activation of NOX2 would mimic the effect of Aβ_1-42_ on glucose utilization. Standard NOX activator PMA (100nM^44^) reproduced the Aβ_1-42_ effects in brain slices, resulting in the reduction of both glucose consumption (Figure S3A,C) and NAD(P)H overshoot (Figure S3B,C). Accordingly, NOX2 antagonist GSK2795039 completely prevented PMA effects (Figure S3B-F).

In the brain, NOX2 is largely expressed in microglia, the resident brain phagocytes, where it is used as a host defense mechanism^26,27^ However, our experiments suggest that neurons^45^ play the dominant role in generating Aβ_1-42_ /NOX2-induced ROS toxicity. While NOX2 in microglia is activated in response to neurotoxic stimulation^25^, NMDA receptor stimulation is required for NOX2 activation in neurons^46^, and we found that NMDAR blockade recapitulated the effect of NOX2 inhibition (Figure S4). Additionally, depleting microglia through dietary PLX5622 supplementation did not ameliorate the Aβ_1-42_ effect on glycolysis (as NAD(P)H transients; Figure S4B) which was again fully prevented by GSK2795039 (Figure S4B), indicating a significant presence of functional NOX2 in our microglia-depleted slices. However, we cannot rule out a possible contribution of microglia as they have also been reported to express NMDARs^47^ (although the presence of functional microglial NMDARs *in situ* is a matter of debate^48,49^) and our microglial depletion experiments did not achieve complete ablation (Figure S4B). Further experiments will also be needed to evaluate a potential contribution of astrocytes which express both NMDARs and NOX.

### NOX2 inhibition prevents Aβ_1-42_-induced hyperexcitability in vivo

Epileptic seizures are a frequent comorbidity of AD^50,51^. Epilepsy also occurs in multiple transgenic mouse models overproducing Aβ^52^, and network hyperactivity has been observed in hippocampal slices following acute Aβ application^53,54^. To investigate the immediate and medium-term effects of Aβ_1-42_ on network electrophysiology, we recorded hippocampal CA1 local field potentials in awake freely-moving mice before and following i.c.v. oligomeric Aβ_1-42_ injection. We observed a rapid onset of network hyperactivity following Aβ_1-42_ injection, seen both as an increased hyperactivity (a general measure of all network activity exceeding 3xSD^55^) and in interictal spike frequency (Figure 3A, C, Table 1.24,25). Network hyperactivity persisted at least up to 48 hours (Figure 3D, Table 1.26,27) after the injection. Furthermore, Aβ_1-42_ induced a significant increase in CA1 pathological high-frequency oscillations (pHFOs; 250-600Hz), a marker for epileptogenesis^56,57^, (Figure 3D, Table 1.28). Importantly, NOX2 inhibition by i.c.v application of GSK2795039 both prior to and following Aβ_1-42_ injection completely prevented all network abnormalities, both acute and 48 hours later (Figure 3B-D, Table 1.29-33). This suggests that NOX2 activation is crucial not only for Aβ_1-42_-induced brain glucose hypometabolism but also for Aβ_1-42_-induced network abnormalities, highlighting the causal link between the two pathologies. It is also in line with our previous reports showing that NOX2 is the primary trigger of epileptic seizures^43^ as well as of brain glucose hypometabolism-induced network hyperactivity and epileptogenesis^35^.

**Figure 3.**
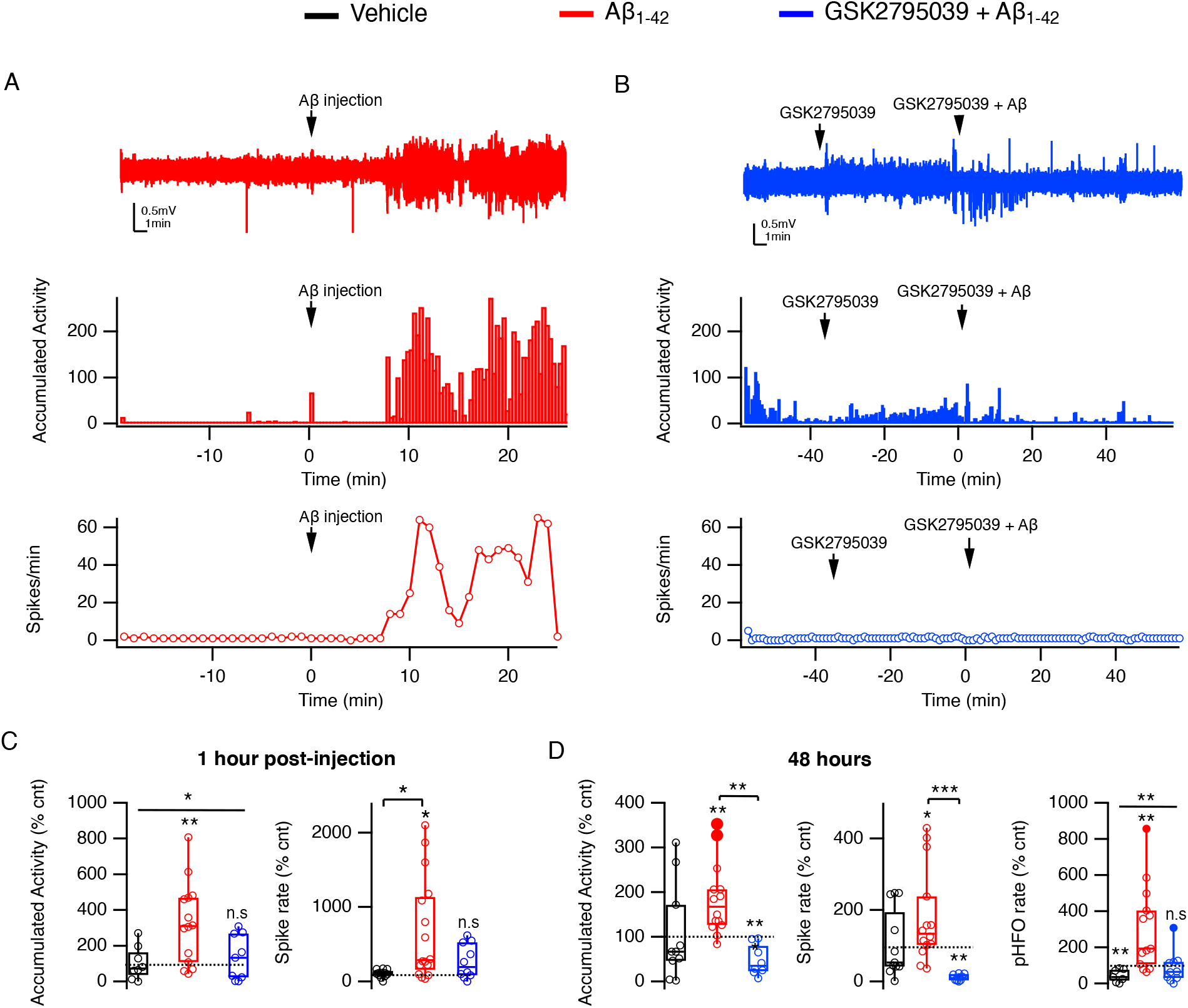
NOX2 mediates Aβ effect on network activity in-vivo. **A.** Aβ_1-42_ i.c.v. injection results in an immediate increase in network activity and interictal spike frequency in awake freely moving mice. Top, example LFP trace from hippocampal CA1 stratum pyramidale before and after Aβ injection. Middle: 20-sec accumulated activity integrals analyzed from the top trace. Bottom: interictal spike frequency analyzed from the top trace. **B.** NOX2 blockade prevents Aβ-induced hyperactivity. Top: example LFP trace from hippocampal CA1 stratum pyramidale in control, following GSK2795039 injection, and after subsequent Aβ injection. Middle: accumulated activity integrals analyzed from the top trace. Bottom: interictal spike frequency analyzed from the top trace. **C.** Mean 1-hour accumulated activity integral and interictal spike rate values for all experiments in A-B. **D.** Aβ-induced hyperactivity persists 48 hours following the Aβ injection and is prevented by preceding GSK2795039 application. Left, average accumulated activity integrals. Middle, average interictal spike rates. Right, average pHFO rates. *p<0.05, **p<0.01, ***p<0.001

### NOX2 inhibition prevents behavioral and psychological symptoms induced by Aβ

We noticed that Aβ-injected mice exhibited clearly increased aggression and anxiety, and evaluated these mouse behaviors 1 week (Open Field and Social Interaction tests) and 2 weeks (Partition Test) after i.c.v. administration of Aβ_1-42_ (Figure 4A). In the Open Field Test^58^, Aβ-injected mice spent twice as long in the central part of the field compared to control mice, signifying elevated anxiety that was further demonstrated by increased defecation (Figure 4B). Mice in the Aβ-group also displayed a much higher level of aggression as revealed by the Social Interaction-Fights test (see Supplementary Video), with the proportion of aggressors as well as the number and total duration of fights increasing sharply (Figure 4C, top). This was paralleled by reduced positive social interactions in Aβ-group: while vehicle-injected mice tended to spend a significant proportion of their time huddled in groups, Aβ-injected mice spent almost no time “cuddling” (Figure 4C, bottom; Supplementary Video). Increased aggression in Aβ-injected mice was also confirmed by the modified Partition Test^59^, where they spent a significantly longer time in direct proximity/contact with the partition separating them from the reference mouse in effort to reach it (Figure 4D). Altogether, these tests demonstrated anxiety and marked aggression in male mice following Aβ_1-42_ injection (see also^60,61^). Importantly, all observed behavioral abnormalities were prevented by daily i.c.v. injections of GSK2795039 (Figure 4B-D), indicating that these neuropsychiatric-like disturbances were caused by NOX2-induced oxidative stress.

**Figure 4.**
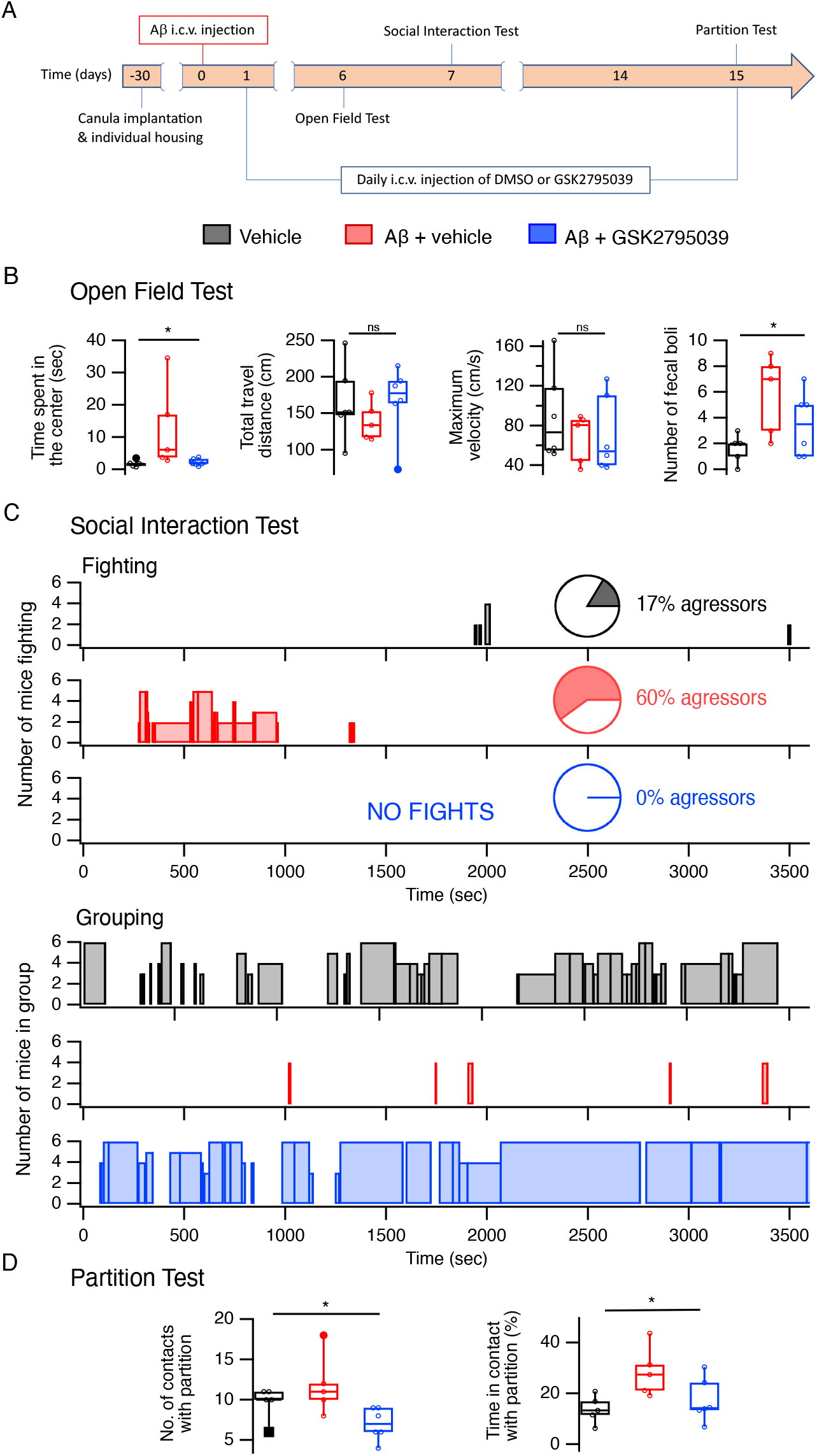
NOX2 blockade prevents Aβ_1-42_-induced neuropsychiatric-like behavioral abnormalities. **A.** Timeline diagram of experimental design. **B. Open field test.** Animals were placed in the center of the OF and latent time (sec) of the first run from the central square of 20 × 20 cm was recorded. Such indicators of anxiety as the number of entries into the center (n), the average time in the center (sec), and the number of fecal boli (n) are also presented. **C. Social Interaction test**: Animals, individually housed for 5 days prior, were placed on a round field for 1 hour. Top panel, Fighting Test: the number of fights, the number of aggressors, and duration of fights are presented. The number of fights and aggressors increased sharply in the Aβ group (red) compared to vehicle group (black). In the Aβ group with daily GSK2795039 injection, not a single fight was recorded during the entire hour of registration (blue). Mice actively interacted, but elements of aggression were absent. Bottom panel, Grouping test: at the same time, direct contact (huddling) between mice was evaluated. Number of mice in a group as well as duration of grouping are presented. Mice in Aβ group (red) spent almost no time grouping, in contrast to the vehicle group (black). GSK2795039 treatment resulted in grouping dynamic improvement to or even over the vehicle levels (blue). *See Supplementary Video for sample recording of all three groups. **D. Partition Test**. As direct contact with the partition correlates with the level of aggressiveness in mice, we analyzed the number of contacts (right) and time in contact with the partition (left). GSK2795039 (blue) prevented Aβ-induced aggressiveness (red) in mice when compared to vehicle (black). *p<0.05.

**Table 1.** Summary of all recorded parameters, presented as % change from controls.

|  | Parameter | Change from control, % | SEM | n | p | Figure |
| --- | --- | --- | --- | --- | --- | --- |
| **Metabolic effects of Aβ** |  |  |  |  |  |  |
| **Aβ** |  |  |  |  |  |  |
| 1 | Extracellular glucose transient amplitude, slices | 57.7 | 4.13 | 21 | < 0.0001 | 1A |
| 2 | Extracellular lactate transient amplitude, slices | 67 | 9.85 | 7 | <0.02 | S1A,B |
| 3 | Extracellular glucose transient amplitude, in-vivo | 248.5 | 45 | 5 | < 0.0005 | 1B |
| 4 | LFP train integral, in-vivo | 101.4 | 7.644 | 5 | 0.8597 | 1B |
| 5 | NAD(P)H "overshoot" amplitude | 56.4 | 3.8 | 23 | < 0.0001 | 1C |
| 6 | NAD(P)H "dip" amplitude | 118.6 | 7.55 | 23 | < 0.03 | 1C |
| 7 | FAD "undershoot" amplitude | 73.1 | 4.32 | 7 | < 0.001 | S1F |
| 8 | pO<sub>2</sub> integral | 118.7 | 6 | 26 | < 0.005 | 1D |
| 9 | LFP train integral, slices | 102.6 | 3.96 | 21 | 0.11 | 1E |
| **GSK2795039 + Aβ** |  |  |  |  |  |  |
| 10 | Extracellular glucose transient amplitude, slices | 109.5 | 10.2 | 7 | 0.39 | 2A |
| 11 | Extracellular glucose transient amplitude, in-vivo | 91.44 | 5.94 | 5 | 0.22 | 2B |
| 12 | LFP train integral, in-vivo | 126.8 | 27.3 | 5 | 0.625 | 2B |
| 13 | NAD(P)H "overshoot" amplitude | 125.2 | 16.47 | 8 | 0.17 | 2C |
| 14 | NAD(P)H "dip" amplitude | 103.6 | 5.841 | 8 | 0.5598 | 2C |
| 15 | pO<sub>2</sub> integral | 100 | 3.677 | 4 | 0.995 | 2D |
| 16 | LFP train integral, slices | 82.09 | 4.711 | 8 | < 0.001 | 2E |
| **Aβ in NOX2-deficient Cybb(tm1din/J) mouse slices** |  |  |  |  |  |  |
| 17 | Extracellular glucose transient amplitude | 88.13 | 6.51 | 5 | 0.1423 | 2F |
| 18 | NAD(P)H "overshoot" amplitude | 93.15 | 11.05 | 10 | >0.9999 | 2F |
| 19 | NAD(P)H "dip" amplitude | 107 | 4.527 | 10 | 0.1572 | 2F |
| 20 | LFP train integral | 114.6 | 7.151 | 18 | 0.0568 | - |
| **Lipid Peroxidation** |  |  |  |  |  |  |
| 21 | MDA, Aβ | 273.2 | 101.4 | 7 | <0.05 | S2 |
| 22 | MDA, GSK2795039 + Aβ | 72.51 | 8.39 | 6 | <0.03 | S2 |
| 23 | MDA, Aβ in Cybb(tm1din/J) mouse slices | 107 | 18.57 | 6 | 0.7201 | S2 |
| **Network activity in freely moving mice** |  |  |  |  |  |  |
| **i.c.v Aβ** |  |  |  |  |  |  |
| 1 hour post-injection |  |  |  |  |  |  |
| 24 | Accumulated Activity | 325 | 60.79 | 14 | <0.003 | 3C |
| 25 | Interictal spike rate | 674.2 | 172.1 | 16 | <0.002 | 3C |
| 48 hours post-injection |  |  |  |  |  |  |
| 26 | Accumulated Activity | 184.3 | 21.55 | 14 | <0.002 | 3D |
| 27 | Interictal spike rate | 179.9 | 35.04 | 14 | <0.05 | 3D |
| 28 | pHFO rate | 304.4 | 66.26 | 13 | <0.01 | 3D |
| **i.c.v GSK2795039 + Aβ** |  |  |  |  |  |  |
| 1 hour post-injection |  |  |  |  |  |  |
| 29 | Accumulated Activity | 134.4 | 41.98 | 9 | 0.4357 | 3C |
| 30 | Interictal spike rate | 269.2 | 73.05 | 10 | 0.0977 | 3C |
| 48 hours post-injection |  |  |  |  |  |  |
| 31 | Accumulated Activity | 47.53 | 12.07 | 8 | <0.004 | 3D |
| 32 | Interictal spike rate | 11.16 | 3.412 | 8 | <0.008 | 3D |
| 33 | pHFO rate | 88.18 | 28.23 | 10 | 0.6853 | 3D |
| **i.c.v Vehicle** |  |  |  |  |  |  |
| 1 hour post-injection |  |  |  |  |  |  |
| 34 | Accumulated Activity | 114.4 | 27.19 | 10 | 0.6092 | 3C |
| 35 | Interictal spike rate | 105.6 | 14.74 | 12 | 0.5557 | 3C |
| 48 hours post-injection |  |  |  |  |  |  |
| 36 | Accumulated Activity | 107 | 33.97 | 10 | 0.8422 | 3D |
| 37 | Interictal spike rate | 107.3 | 29.06 | 11 | 0.9502 | 3D |
| 38 | pHFO rate | 43.65 | 13.3 | 7 | <0.006 | 3D |

## Discussion

Glucose hypometabolism is implicated in the initiation of sporadic AD, as it is associated with many major AD risk factors^10–13^. It occurs in patients with amnestic mild cognitive impairment (aMCI), widely thought to be a prodromal stage of AD^10,14,15^, and has also been detected in AD patients almost two decades prior to the onset of clinical symptoms. Importantly, brain hypometabolism in AD is always associated with oxidative stress (for review,^12,15^). Energy deficiency^62,63^ and oxidative stress^64^ have both been reported to result in BACE1 upregulation and increased production of Aβ. The resulting accumulation of Aβ_1-42_ is key to the pathogenesis of AD and is known to induce oxidative stress^17^. We now show that Aβ inhibits brain glucose utilization by inducing NOX2-mediated oxidative stress, thus establishing a vicious cycle of AD pathogenesis. While the correlation between the Aβ load and NOX2 activity has been reported in multiple studies^19,28–33^, until now functional consequences of this interaction were unclear. Downstream of NOX-2 activation with the resulting brain hypometabolism, our results show that these alterations lead to network hyperactivity and neuropsychiatric-like changes in animal behavior, establishing a causal chain between multiple AD-related pathologies.

Importantly, our data suggests that Aβ-driven disturbance of glucose metabolism is restricted mainly to cytosolic (aerobic) glycolysis while oxidative phosphorylation is actually upregulated, presumably as a compensatory mechanism as has been suggested by previous studies^37,65^ (also^66, 38^). Our observation may seem out of line with established dogma since metabolic changes in AD brain are thought to reflect or include impaired mitochondrial function^67^. However, the underproduction of mitochondrial ATP may be the consequence of either insufficient support provided by glycolysis and/or impaired process of oxidative phosphorylation including the TCA cycle. Early studies reported that early in AD, the cerebral metabolic rate of oxygen was not altered or was changed disproportionally to the prominent decrease in glucose utilization^68–70^. It was hypothesized that unaltered oxygen utilization and normal CO_2_ production may indicate undisturbed substrate oxidation in mitochondria^68^. Moreover, other early studies that used the arterio-venous difference method showed that brain ketone uptake is still normal in moderately advanced AD^71,72^, while ketone catabolism is entirely mitochondrial. Studies using PET ketone tracer 11C-acetoacetate (AcAc) reported that brain metabolism of ketones is unchanged in MCI and early AD^73,74^ supporting the previous assumption that oxidative phosphorylation may still be normal in early AD. Finally, a recent study utilizing RNA-seq on *post mortem* AD brains showed impaired glycolytic pathways in neurons and astrocytes while the ketolytic pathways remained normal^75^. Altogether, this suggests that early brain hypometabolism in AD may be specific to glycolytic breakdown^76^ and not in dysfunctional mitochondrial oxidative phosphorylation.

The NOX family of enzymes are transmembrane proteins that transport an electron from cytosolic NADPH to reduce oxygen to superoxide anion. There are seven known isoforms of NOX, with NOX1, NOX2 and NOX4 expressed in multiple brain regions including the cerebral cortex, hippocampus, cerebellum, hypothalamus, midbrain and/or striatum^23^. Of these NOX variants, NOX2 is the predominant form expressed by microglia, neurons, and astrocytes. Several lines of evidence including postmortem analyses of AD patient cerebral cortices indicate that oxidative stress—particularly resulting from NOX2 activation—plays a significant role in the development of AD^20,23,25–27^. The close relationship between the levels of Aβ and NOX2 activity has also been well documented in multiple studies^19,28–33^.

The selective NOX2 antagonist GSK2795039 that we used in our study is a novel small-molecule NOX2 inhibitor that is a first of its kind with brain bioavailability following systematic (oral) administration. GSK2795039 is not cytotoxic at concentrations effective for NOX2 inhibition^77^: it was well tolerated in rodents, with no obvious adverse effects following 5 days of twice-daily dosing^77^. Following oral administration, GSK2795039 was detected in the blood and the central nervous system, indicating that it can cross the blood-brain barrier^77^. Thus, to our knowledge GSK2795039 is the only currently available selective NOX2 inhibitor that may be appropriate for the development of future AD therapies^78^.

Given the primary role of oxidative stress in AD pathology, one other alternative treatment strategy could be scavenging ROS by antioxidants. However, clinical trials in aMCI and AD patients using antioxidants have so far been disappointing^12,79^. Our previous studies might provide an explanation for this apparent discrepancy as we showed that the potent antioxidant Tempol^80,81^ failed to abate rapid ROS accumulation resulting from NOX activation during epileptic seizures^43^. It thus appears necessary to instead go to the source and to block pathological NOX2 activation directly in order to prevent its detrimental effects.

Intriguingly, oxidative stress (particularly NOX-induced oxidative stress), neuroinflammation, and hypometabolism – all acting in concert - have been reported in the early stages of other major neurodegenerative diseases^82–85,20,86^. Therefore, we posit that the role of NOX-hypometabolism-network dysfunction axis is pervasive in neurodegenerative disease initiation, and special attention should be paid to NOX2 in search of effective treatments.

In summary, in this study we established direct causal links between Aβ_1-42_, NOX2-induced oxidative stress, glucose hypometabolism, network hyperactivity, and neuropsychiatric disturbances in AD pathogenesis. We also demonstrate the potency of selective NOX2 inhibitor GSK2795039 in preventing toxic Aβ_1-42_ effects and propose NOX2 as a primary target for early interventions in Alzheimer’s disease, warranting further studies.

## Materials and Methods

All animal protocols and experimental procedures were approved by the Ethics Committees for Animal Experimentation at the INSERM (#30-03102012), ITEB RAS, the University of California and Gladstone Institutes under IACUC protocol AN176773.

### Experimental animals

Experiments were performed on mature (2-4 months) male mice. A number of strains was utilized: OF1 (Charles River Laboratories, France), C57/Bl6 mice (Jackson Labs, USA), or BALB/c mice (Laboratory Animal Nursery “Pushchino”, Pushchino, Russia).

### Ex vivo experiments

#### Tissue slice preparation

Mice were anaesthetized with isoflurane and decapitated; the brain was rapidly removed from the skull and placed in the ice-cold ACSF. The ACSF solution consisted of (in mmol/L): NaCl 126, KCl 3.50, NaH_2_PO_4_ 1.25, NaHCO_3_ 25, CaCl_2_ 2.00, MgCl_2_ 1.30, and dextrose 5, pH 7.4. ACSF was aerated with 95% O_2_/5% CO_2_ gas mixture. Sagittal slices (350 μm) were cut using a tissue slicer (Leica VT 1200s, Leica Microsystem, Germany). During cutting, slices were submerged in an icecold (< 6°C) solution consisting of (in mmol/L): K-gluconate 140, HEPES 10, Na-gluconate 15, EGTA 0.2, NaCl 4, pH adjusted to 7.2 with KOH. Slices were immediately transferred to a multisection, dual-side perfusion holding chamber with constantly circulating ACSF and allowed to recover for 2h at room temperature (22°C-24°C).

#### Synaptic stimulation and field potential recordings

Slices were transferred to a recording chamber continuously superfused (10 ml/min) with ACSF (33-34°C) with access to both slice sides. Schaffer collaterals/commissures were stimulated using the DS2A isolated stimulator (Digitimer Ltd, UK) with a bipolar metal electrode. LFPs were recorded using glass microelectrodes filled with ASCF, placed in stratum pyramidale of CA1 area and connected to ISO DAM-8A amplifier (WPI, FL) or MultiClamp 700B amplifier (Axon Instruments, USA). Synaptic stimulation consisted of a 30-second stimulus train (300 pulses) at 10Hz. Stimulus current was adjusted using single pulses (40-170μA, 200μs) to induce a LFP with a population spike of about 50% maximal amplitude.

#### Oxygen, glucose and lactate measurements

A Clark-style oxygen microelectrode (Unisense Ltd, Denmark) was used to measure slice tissue partial oxygen pressure (pO_2_). Tissue glucose and lactate concentrations were measured with enzymatic microelectrodes (tip diameter 25μm; Sarissa Biomedical, Coventry, UK) connected to a free radical analyzer TBR4100 (Word Precision Instruments Ltd, UK). Calibration using known multiple substrate concentrations was performed after the first polarization and was repeated following each experiment to ensure the sensor’s unchanged sensitivity to a substrate.

#### NAD(P)H and FAD fluorescence imaging

Changes in NAD(P)H and FAD autofluorescence in hippocampal slices were recorded as described previously^36^. Data were expressed as the percentage change in fluorescence over baseline [(DF/F)·100].

#### Pharmacology

Antagonist of NMDA receptors, (2R)-amino-5-phosphonopentanoate (APV) was purchased from Tocris Bioscience (Bio-Techne Ltd, UK); GSK2795039 from MedChemExpress.

#### Aβ_1-42_ preparation

Oligomeric Aβ_1-42_ (Sigma Aldrich, USA) was prepared according to manufacturer specifications. Briefly, Aβ_1-42_ was dissolved in 0.2% NH_4_OH at 1mg/mL, brought to 400nM concentration in ACSF and sonicated for 1min. Prior to experiments, solution was allowed to fibrillize for minimum of 1 hour.

#### Lipid Peroxidation Assays

Acute sagittal brain slices were prepared as described in the previous section and were allowed to recover in a vapor interface holding chamber (Scientific Systems Inc, Canada) containing standard ACSF at 34C for one hour. Slices were then assigned in an alternating pattern to either the control or the Aβ group, so that each slice incubated in Aβ-containing ACSF had its immediate neighbor from same hemisphere incubated in control ACSF. Following 1-hour incubation in their respective oxygenated ACSF solutions, slices were snap-frozen over dry ice. Slices were then homogenized using a 26-gauge needle in the presence of a high-detergent buffer consisting of 50 mM Tris, 150 mM sodium chloride, 2% Nonidet P-40, 1% sodium deoxycholate, 4% sodium dodecyl sulfate, and supplemented with complete protease inhibitor cocktail (Roche), phosphatase inhibitor cocktail 1 (P2850, Sigma), phosphatase inhibitor cocktail 2 (P5726, Sigma), and 0.005% Butylated hydroxytoluene. The total protein in cell lysates was quantitated with the BCA protein assay kit (#23227, Pierce). Malondialdehyde (MDA) levels were quantified using a OxiSelect™ MDA Adduct Competitive ELISA Kit (Cell Biolabs, Inc, USA) as per manufacturer instructions. Recorded MDA levels were normalized to each sample’s protein concentration and the result then normalized to the average of all control slices from same hemisphere.

### In vivo electrophysiology on freely-moving animals

#### Animals and surgery

Before the experiment, mice were implanted with nichrome recording electrodes in the CA1 (AP=-2.5mm, ML= 2mm, H=1.5mm) and dentate gyrus (AP=-2mm, ML= 0.8mm, H=2.5mm), and a guide cannula for i.c.v. injections (AP=-0.2mm, ML= 1.8mm, H=2.2mm). Animals were anesthetized with Zoletil (120 mg/kg) supplemented with xylazine (10mg/kg). Electrode and cannula placements were verified post-mortem. A reference electrode was screwed into the occipital bone.

#### Drug administration

Animals received either 1μl vehicle (NaCl 0.9%+PBS+Glucose 5mM) or 1μl of Aβ_1-42_ (Sigma-Aldrich, 4mg/ml in 1.0% NH_4_OH) intracerebroventricularly (i.c.v.) using a Hamilton syringe. In experiments with GSK2795039, 1μl of inhibitor solution (20mg/ml) was injected 30 min prior to Aβ_1-42_ and additionally 1μl together with Aβ_1-42_ solution. After a four-day recovery period, LFP recordings on freely-moving animals were initiated. Following 60-minute recordings of control activity, animals received an injection of the appropriate drug. To estimate acute effects, brain activity was monitored 90 min after the drug application. For 48 hours following the experiment, animals received daily i.c.v. injection of either vehicle (control and Aβ groups) or GSK2795039 (Aβ+ GSK2795039 group). 1-hour long LFP recordings were performed 48h following the acute experiments.

#### LFP recordings

LFPs were filtered (high-pass filter 0.1Hz, low-pass filter 5kHz) and recorded at 10kHz sampling rate. Accumulated activity integrals and interictal spikes were detected as described previously^35,55^ using custom macros in IgorPro (Wavemetrics, USA). Briefly, accumulated activity constitutes a sum of all events exceeding the 3xSD threshold and binned at 20sec intervals. Interictal spikes were detected as any event exceeding 5xSD amplitude and lasting between 10-90msec. Fast ripples were detected as events with Hilbert transform envelope exceeding 3xSD threshold, minimum of 3 oscillations, and mean frequency between 250-600Hz.

### In vivo glucose experiments on anesthetized mice

#### Animals and surgery

For acute glucose measurements, 10 mice (Aβ group, n=5, Aβ+GSK2795039 group, n=5) were anesthetized with pentobarbital (60mg/kg) supplemented with xylazine (20 mg/kg). Animals were placed in a stereotaxic frame, scalped, and holes for electrodes and guide cannula were drilled. In addition to hippocampal field electrode and guide cannula, a bipolar stimulating electrode was implanted to the fimbria fornix (AP=-0.4mm, L= 1mm, H=3mm, approach angle 15°) for hippocampal network activation. A cranial window (Ø 1.5mm) above the hippocampus was drilled ipsilateral to the cannula, and a Sarissa glucose sensor was dipped in the hippocampus (2mm) using a micromanipulator. After surgical preparation, the cortex was kept under saline to prevent drying.

#### LFPs and extracellular glucose measurements

LFPs and extracellular glucose were measured using the same equipment as for the *in vitro* experiments, with network activity induced by 10Hz, 30sec stimulation of Schaeffer collaterals and LFPs and glucose measured in CA1 region. Following 2-3 stimulations (15 min apart) in control conditions, Aβ_1-42_ or GSK2795039 followed by GSK2795039 + Aβ_1-42_ were injected, with 2-3 recordings performed in each subsequent condition.

#### PLX treatment for microglial depletion

To deplete microglia, mice were switched to PLX5622 diet (PLX5622-enriched chow was provided by Research Diets, Inc., New Brunswick, USA). Mice were on the ad libitum diet for 7-15 days. Approximately 5 mg of PLX5622 was ingested daily by mice of 35 grams of weight.

#### Histological analysis

Free-floating slices (35 μm) from perfused animals were incubated with Triton X (0.3%) in PBS three times for 5 min, followed by blocking solution (BSA 1%, Triton X 0.3% in PBS) for 2 h. Then, the primary antibody rabbit anti-Iba-1 (1:1000; Wako, Japan) was added and slices were incubated overnight at 4°C. Next day, the secondary antibodies goat anti-rabbit (1:1000; Alexa Fluor 488, ThermoFisher, USA) were added for 2 h. After washout in PBS with 0.3% TritonX-100, slices were mounted on gelatinized covers in Fluoromount media (Sigma-Aldrich, USA). Immunostaining was analyzed under a Nikon E200 fluorescence microscope. In order to make a proper comparison, equivalent regions containing similar portions were chosen for all the groups. 3+ sections per animal were used for averaging. Photomicrographs using 10X (0.25 of numerical aperture) of stained fluorescence were quantified with the aid of ImageJ software (NIH, USA), and the whole hippocampus was used for quantification. The number of Iba-1+ cells was calculated using custom macros.

### Behavioral tests on effects of Aβ administration

Male mice were housed individually with food and water ad libitum. Animals were maintained with a 12-h light/dark cycle (lights on from 9 AM to 9 PM) in temperature (22°C ± 1°C)-controlled room. The protocol was approved by the Committee on the Bioethics of Animal Experiments of the Institute of Theoretical and Experimental Biophysics of the Russian Academy of Sciences.

#### Surgery

Animals underwent a neurosurgical operation under general anesthesia: mixture ketamine (200 mg/kg, i.m.) and Xylazine hydrochloride (10 mg/kg, i.p.). Premedication with atropine (0.04 mg/kg) was performed subcutaneously 15 minutes before surgery. The body temperature was maintained by a heating pad and the cardiopulmonary parameters were monitored during the surgery by an Oxy9Vet Plus pulse oximeter (Bionet, South Korea). Guide cannula (stainless steel, 21 gauge) was implanted above the left lateral brain ventricle (AP = −0.7; L = 1.37; H = 1.25) according to the mice brain atlas^87^.

#### Drug administration

Drugs were administered 1-month post-surgery to avoid surgery side effects. Mice were allocated to three groups: control (n = 6), Aβ (n = 5), and Aβ+GSK2795039 (n = 6). Injections were made i.c.v. through the guide cannula (1 μl/min) to awake mice. All substances were injected in equal volumes (1 μl) and the guide cannula was closed by a plunger. Mice in the Aβ+GSK2795039 group received GSK2795039 (MedChemExpress Europe; 4.5 mg/ml in DMSO) and oligomeric Aβ_1-42_ (Sigma-Aldrich, 1 mg/ml in 1.0% NH_4_OH) on the first day, and daily GSK2795039 further for 14 days. Mice in the Aβ group received Aβ_1-42_ on the first day and daily DMSO for the following 14 days. Mice in the control group received daily DMSO for 15 days.

#### Behavioral assessment

Experiments were carried between 18:00 and 21:00. The EthoVision system (Noldus Information Technology, Wageningen, the Netherlands) and RealTimer (Open Science, Moscow, Russia) were used for video registration and subsequent analysis.

#### Open field test

To evaluate anxiety, the animals’ behavior was observed in an “open field”, which was a square, black area 60 × 60 cm in size with sides 40 cm high^58^. The central area was defined as a 20 cm × 20 cm square, and the other region was defined as the peripheral area. Animals were placed initially in the center of the “field”. The time spent in the field center, total travel distance and maximal velocity were recorded and analyzed.

#### Social Interaction Test

Social interaction tests were used on mice accustomed for 5 weeks to social isolation. To evaluate aggressive interaction in a group of animals the “fights” evaluation was used. Animals of the same experimental group were placed in the round field (diameter 50 cm) for 1 hour. Each mouse was marked with a colored spot on its back for identification. Aggressive behavior estimated as number of attack defined as rushing and leaping at a partner with bites and kicks (in accordance with^88^) (Supplementary Video, see for example from 10:05 in Ab group). The number of episodes of fights, the average duration of fights, the number of participants in fights, and the number of “aggressors” (mice who initiated the fight) were analyzed. Positive social interaction in groups of mice was assessed by “grouping” episodes (Supplementary Video, see for example from 00:16 in control + GSK groups). Such social interaction was defined as a close and friendly grouping of 3 or more mice for at least 4 seconds. To estimate this type of behavior, number of grouping episodes, average duration and number of participants were analyzed.

#### Partition test

The Partition test directly correlates with the level of aggressiveness in mice^59^. An experimental mouse was placed in a large compartment (24.1 × 17.8 × 14.0 cm) and a reference mouse in a smaller compartment (6.4 × 17.8 × 14.0 cm), with compartments separated by a see-through partition. For 5 minutes of the test, the number of approaches to the partition and the total time spent near the partition (when the mouse touches it with its paws or nose, reacting to a partner in the neighboring compartment), were recorded and analyzed.

### Statistical analysis

Normality was determined by the Kolmogorov-Smirnov test. Significance was determined by the student t-test or one-way ANOVA with Tukey’s or Kruskal-Wallis multiple comparisons post-hoc tests, where appropriate. Data are presented as mean ± SEM.

## List of Supplementary Materials

Supplementary Figure 1, Supplementary Figure 2, Supplementary Figure 3, Supplementary Video.

## Acknowledgments

This study was supported by the National Institute on Aging grant R01AG061150 to MZ, RSF grants #17-75-20245 to AM and #20-65-46035 to IP and AO. This work was also supported by NIH/NCRR grant C06 RR018928 to Gladstone Institutes. We thank Dr. Kathryn Claiborn for editorial assistance.

## Author contributions

AM, AI, IP, AO, MZ, and SSJ carried out the study. MZ and YZ designed and coordinated the study, supervised the project, analyzed data, and wrote the manuscript. S.Y.Y. managed mouse lines. Y.H. provided advice on study design and critically reviewed the manuscript.

## Data and materials availability

All data associated with this study are available in the main text or the supplementary materials.

