## Supplemental Figures for "Aβ initiates brain hypometabolism and network dysfunction via NOX2 activation: a potential onset mechanism of Alzheimer’s disease"

Supplementary Figure 1

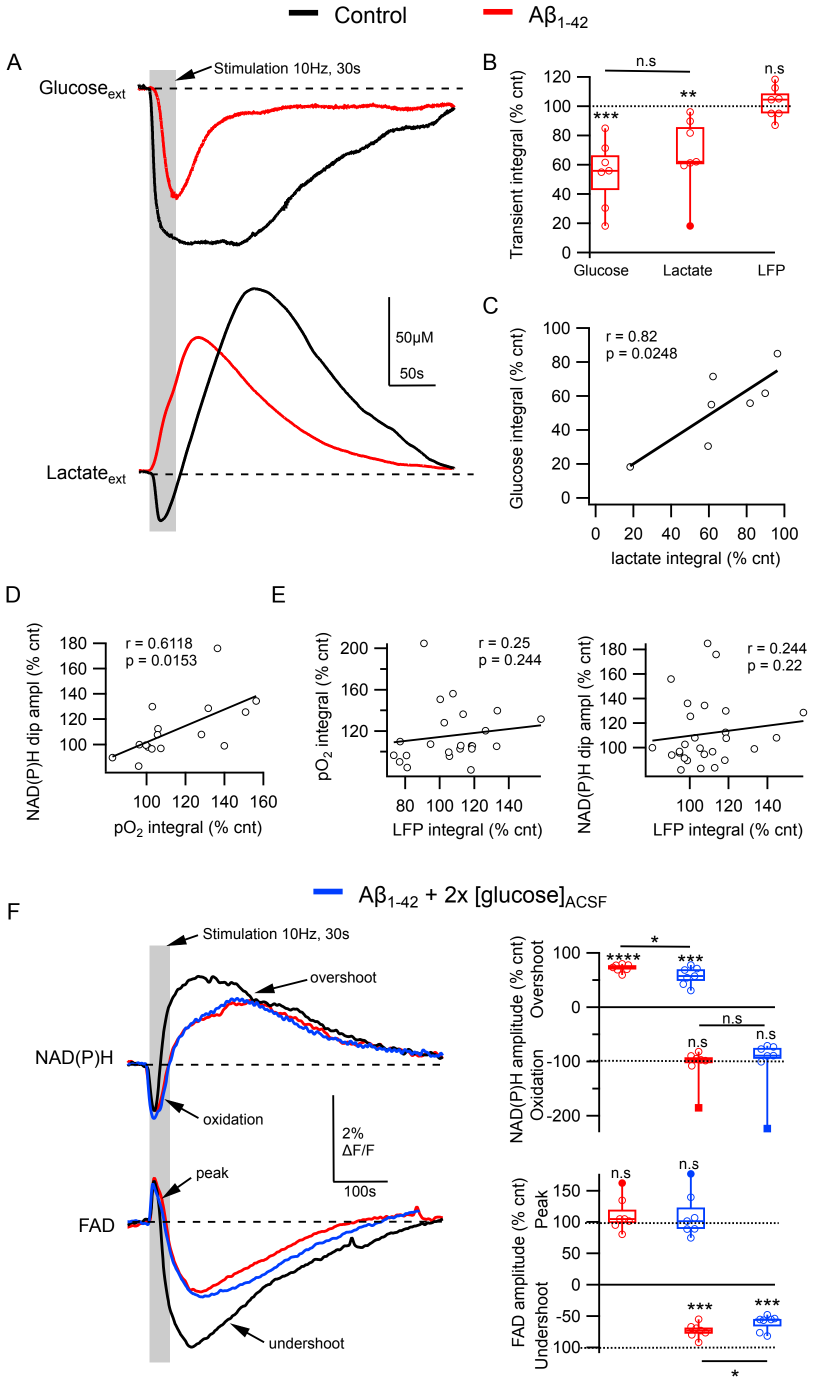


**Figure S1.** ***Multiple effects of Aβ toxicity on glucose metabolism and glycolysis.*** **A**. Example traces from a single experiment in brain slice with activity-driven extracellular glucose and lactate transients measured simultaneously in CA1 in control (black) and following 40 minutes of $A\beta_{1-42}$ application (red). **B**. Summary of all such experiments. **C**. Decrease in activity-driven lactate release is significantly correlated with reduction in glucose uptake under Aβ. **D.** Aβ-induced increase in NAD(P)H oxidation phase amplitude is positively correlated with the parallel increase in oxygen consumption. **E.** Changes in metabolic transients are not correlated with changes in network response seen as LFP integrals. **F.** Increasing extracellular glucose concentration has no effect on Aβ-induced disruption of glycolysis. Left, Example traces from a single experiment in brain slices with activity-driven NAD(P)H and FAD fluorescence transients measured simultaneously in control (black), following 40 minutes of $A\beta_{1-42}$ application (red), and after subsequent doubling of ACSF glucose concentration (blue). Note that Aβ-induced reduction in NAD(P)H overshoot is paralleled by a reduction in FAD "undershoot", confirming inhibited glycolysis. Right, summary of all such experiments. *$p < 0.05$, **$p<0.01$, ***$p<0.001$, ****$p<0.0001$.

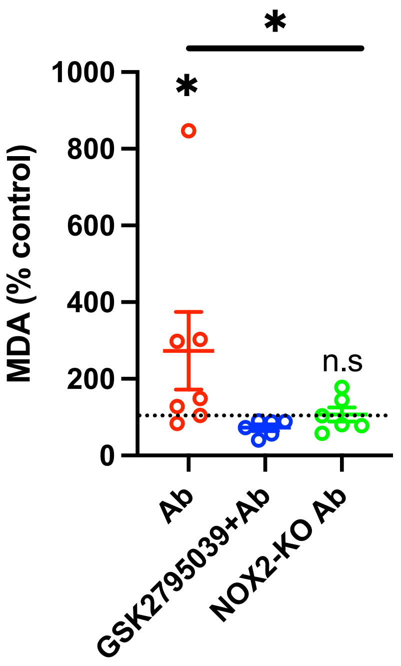


**Figure S2.** ***Aβ induces lipid peroxidation via activation of NOX2.*** Individual brain slice MDA levels, normalized to corresponding non-Aβ controls. Red: WT slices exposed to 400nM fibrillar $A\beta_{1-42}$ for one hour; Blue: WT co-treated with $A\beta_{1-42}$ and GSK2795039; and Green: Brain slices from NOX2-deficient ($Cybb^{tm1din}$/J) mice exposed to $A\beta_{1-42}$. $*p<0.05$

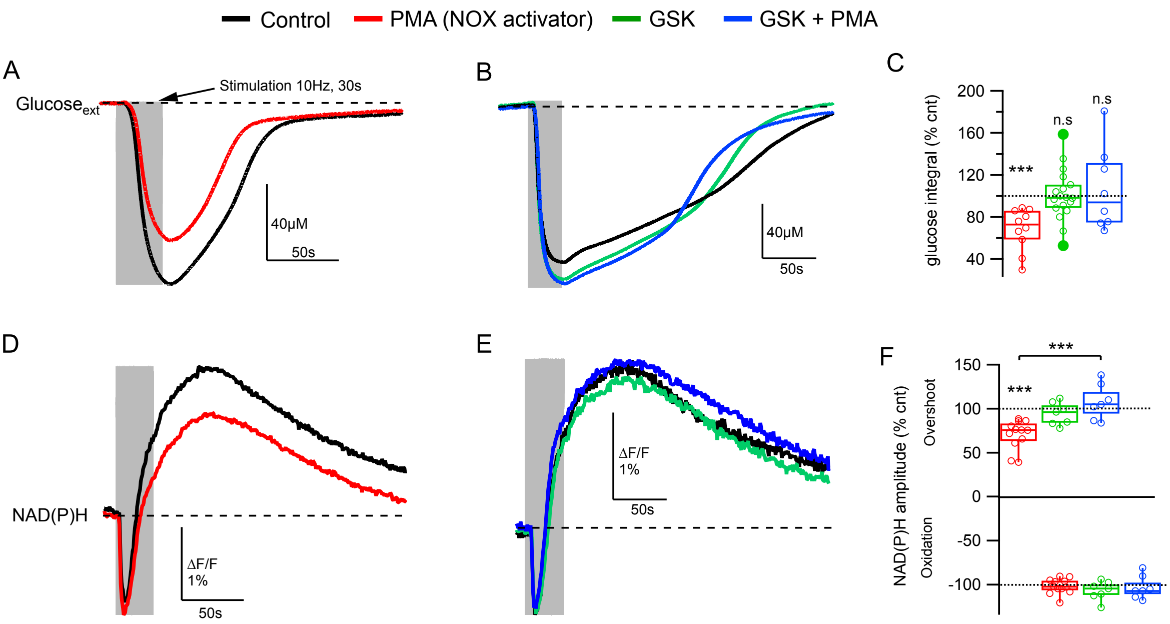


**Figure S3.** ***Direct activation of NOX by PMA reproduces the effect of*** **A**$\beta_{1\text{-}42}$ ***on glucose utilization***. **A.** Application of NOX activator PMA (100nM) reduces glucose uptake; average traces from a single experiment showing extracellular glucose transients in CA1 pyramidal cell layer in response to a 10Hz, 30s stimulation of Schaffer collaterals (grey) in control (black) and following 40min of PMA application (red) **B.** GSK2795039 prevents the effects of PMA on activity-driven glycolysis. Example traces from a single experiment showing glucose transients in control (black), after application of GSK2795039 (green), and following GSK2795039+PMA application (blue). **C**. Summary graph of all corresponding glucose transient integral values in A-B. **D**. PMA reduces glycolysis: NAD(P)H autofluorescence traces from a single experiment in control (black) and after application of PMA (red). **E**. PMA effect on glycolysis is blocked by GSK2795039: NAD(P)H autofluorescence traces from a single experiment in control (black), after application of GSK2795039 (green), and following GSK2795039+PMA application (blue). **F**. Summary graph of all corresponding NAD(P)H amplitudes in D-E. ***p<0.001

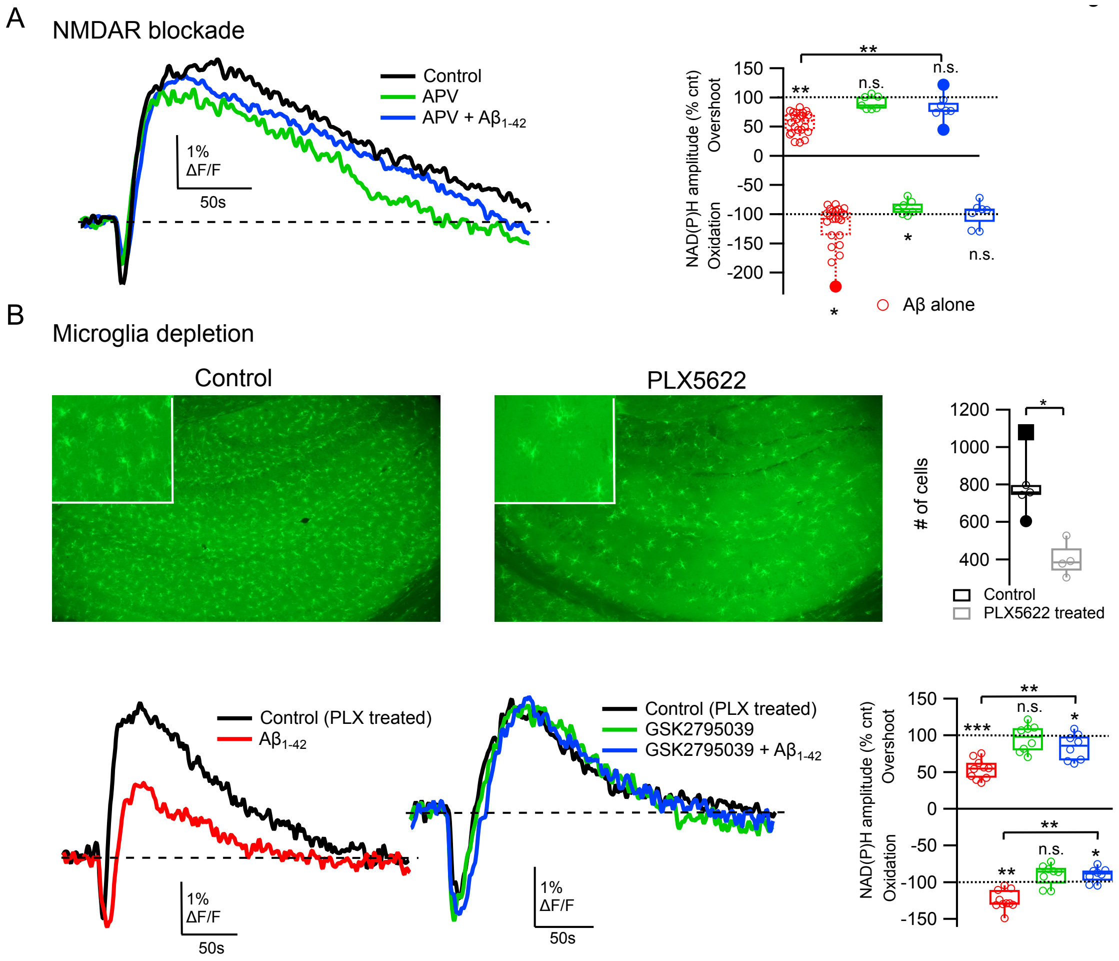


**Figure S4.** ***Locus of NOX2 expression behind Aβ-induced glucose hypometabolism.***
**A.** NMDA receptor blockade by APV prevents Aβ effects on glycolysis, suggesting neuronal NOX contribution. Left, example traces from a single experiment showing activity-induced NAD(P)H autofluorescence transients in control (black), following APV application (green), and after subsequent application of $A\beta_{1\text{-}42}$ (blue). Right, the summary plot of NAD(P)H oxidation and overshoot amplitudes; red markers indicate Aβ values from Fig. 1 for comparison. **B, C.** Microglial depletion does not prevent Aβ effect on glucose utilization. **B.** Sample images of *ex-vivo* hippocampal microsections stained for Iba1 antibody from mice following 1 week of control (left) and PLX5622-containing diet (center). Right, Iba1+ cell counts across all slices. **C.** Hippocampal slices from PLX5622-treated mice are not protected from Aβ toxicity on glucose utilization. Left, traces from a single experiment showing activity-induced NAD(P)H autofluorescence transients in control (black) and following application of $A\beta_{1\text{-}42}$ (red). Center: NOX2 blockade in microglia-depleted slices prevents Aβ effects on glycolysis. Right: summary plot of corresponding NAD(P)H amplitudes. *p<0.05, **p<0.01, ***p<0.001
